# Fitting Force Field parameters to NMR Relaxation Data

**DOI:** 10.1101/2023.02.10.527984

**Authors:** Felix Kümmerer, Simone Orioli, Kresten Lindorff-Larsen

## Abstract

We present an approach to optimise force field parameters using time-dependent data from NMR relaxation experiments. To do so, we scan parameters in the dihedral angle potential energy terms describing the rotation of the methyl groups in proteins, and compare NMR relaxation rates calculated from molecular dynamics simulations with the modified force fields to deuterium relaxation measurements of T4 lysozyme. We find that a small modification of C^*γ*^ methyl groups improves the agreement with experiments both for the protein used to optimize the force field, and when validating using simulations of CI2 and ubiquitin. We also show that these improvements enable a more effective *a posteriori* reweighting of the MD trajectories. The resulting force field thus enables more direct comparison between simulations and side-chain NMR relaxation data, and makes it possible to construct ensembles that better represent the dynamics of proteins in solution.

## 1 Introduction

Understanding the structure and dynamics of proteins is key to determine and manipulate the function of proteins. Experimentally, nuclear magnetic resonance (NMR) spectroscopy is extensively used in both protein structure determination and for the investigation of dynamics on a wide range of time scales. Protein side-chain dynamics that occurs on a timescale faster than the overall rotational motion (i.e. the picosecond to low nanosecond timescale) can play a critical role in a protein’s function by contributing to the conformational entropy.^1–11^ Such dynamics is typically probed by NMR using spin relaxation experiments^12–15^ and although relaxation parameters are very informative, the intricate nature of spin relaxation often makes it difficult to interpret the measurements and distinguish contributing processes. One way around this is to combine NMR experiments with molecular dynamics (MD) simulations. MD simulation is another key technique to study protein structure and motions and makes it possible to obtain a physical description of such dynamics at atomistic resolution.

The integration of NMR and MD has proven to be particularly useful both to interpret experiments^16–20^ but also for the validation and parameterisation of force fields.^21–25^ The underlying mechanisms of NMR spin relaxation experiments are largely understood^13^ and thus relatively good models exist to calculate relaxation rates from MD simulations for both the backbone amides and side chains.^16,17,26–30^ In the case of side chain relaxation, it has, however, previously been shown that commonly used force fields did not fully capture the relevant motions or timescales.^29,31–33^ In particular, it was demonstrated that calculated and experimental deuterium relaxation rates for the fast dynamics of methyl-bearing amino acids did not match within experimental error. Hoffman and co-workers suggested that this was in part due to too high methyl rotation barriers, and reparameterised the corresponding force constants of methyl-bearing amino-acids using CCSD(T) coupled cluster quantum chemical calculations of isolated dipeptides. This led to a substantial improvement in the agreement of calculated and experimental NMR relaxation rates and thus the predictive power of the respective force fields. Later work suggested, however, that there are remaining discrepancies, sometimes resulting in underestimated calculated relaxation rates, especially for methyl groups in valine residues.^34^ As a consequence, we here seek to further improve the force field reparameterisation previously published by Hoffmann et al.^29,31^

Generally, there are at least two different approaches to parameterise a force field for MD.^35–37^ One common strategy is to use quantum mechanical (QM) calculations to help obtain values for force field parameters such as torsion potentials.^29,38–41^ While QM-based approaches can provide very accurate results, they are limited by the accessible system sizes and the number of configurations that may be studied due to high computational costs. In the case of protein force fields, QM-based parameterisation is therefore generally performed on smaller model systems such as dipeptides. Some effects that come with larger systems such as side-chain packing are thus not necessarily taken into account.^42,43^ The second approach is to fit force field parameters to match experimental data.^25,44–48^ Naturally, this is limited by the amount of available experimental data and this can, ultimately, lead to both underdetermination or overfitting of the data. Another critical point is the availability of good and reliable models to calculate experimental observables. However, the use of experiments has the benefit that they provide an accurate representation of real-world systems, including a wide range of conditions and larger, biologically more relevant, setups. Often, a hybrid strategy is chosen which can be done by fitting the force field to a combination of QM calculations and experimental data, with the goal of achieving the best overall agreement between the two sets of data.^40,49–52^ In that case, experimental data also proves to be particularly useful for validation of force fields. In recent years, machine-learning-based approaches to derive force field potentials have become more and more relevant with the aim to reach the accuracy of *ab initio* calculations at a lower computational cost. ^53–56^ Although still often focused on smaller systems bridging the gap between QM-based approaches and classical force fields, machine learned force fields for proteins are likely to play an increased role in the future.^57,58^

Here, we use the improved, QM-based force field by Hoffmann et al.^29^ (which we here term HMS) as a starting point for further parameterisation of methyl spinning barriers. We first benchmark the force field by comparison to NMR relaxation data, and then directly integrate the experimental NMR relaxation rates in the optimisation process. Our approach is inspired by previous work of Aliev and co-workers,^48^ who optimised torsional parameters of prolines by fitting NMR scalar (*J*) couplings and ^13^C relaxation rates using MD simulations.

Similar to their approach, we scan a range of relevant parameters and match calculated relaxation rates and NMR data. Specifically, we perform MD simulations of T4 lysozyme (T4L) using a series of force field parameters, calculate relaxation rates via a spectral density mapping approach^16,17,30,34^ and compare them to deuterium spin relaxation rates obtained from NMR experiments. We find that fine tuning the methyl rotation barrier in valine and isoleucine residues is sufficient to improve further the match between experiments and simulations and validate our parameter set, 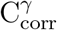, using the model proteins ubiquitin and CI2.

## 2 Results and Discussion

### 2.1 Selecting methyl group rotation parameters

Previous work by Hoffmann and co-workers focused on the reparameterisation of methyl group rotation barriers using CCSD(T) calculations of isolated dipeptides and resulted in substantial improvements—compared to the original, unmodified force fields—when calculating NMR relaxation data.^29,31^ Our subsequent study on the development of a reweighting approach for NMR relaxation parameters revealed, however, remaining issues with the calculated relaxation rates of some residues.^34^ We showed that this was especially problematic in the case of C^*γ*^*H*^*γ*^-bearing side-chains such as VAL and ILE for which the experimental NMR relaxation rates were usually underestimated, implying that, for those methyl groups, the parameters describing methyl rotation might be improved further.^11,34^

We therefore set out to further optimise the force constants of the dihedral angle potential energy terms describing the rotation of the methyl groups (*k*_*dih*_) in the a99SB*-ILDN force field^38,39,44b^ using the HMS corrections by Hoffmann et al, ^59^ 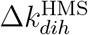, as a starting point for our parameter search (Table 1) for the force constants for the different methyl groups (Table. 2). To do so, we scanned different Δ*k*_*dih*_ values starting from the HMS force field by step-wise increasing Δ*k*_*dih*_ and thus moving towards the original, unmodified force field (Table 3). Additionally, we lowered *k*_*dih*_ for two more steps, and thus examined a total of 15 sets of parameters (i.e. force fields). For each of these force fields, we ran five 1 µs long simulations of T4 lysozyme (T4L) (Table 4), calculated the three NMR relaxation rates *R*(*D*_*z*_), *R*(*D*_*y*_), and 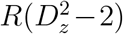 (Eq. 5-7) and compared them to experimentally measured data at 950 MHz^30^ in order to select an optimised parameter set (see Methods section for further details).

**Table 1:**
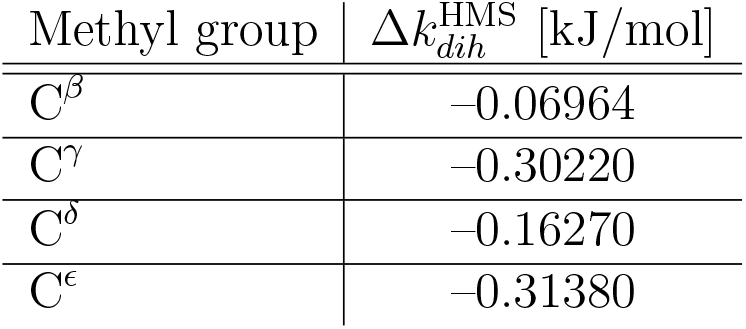
Methyl-group-specific changes to the force constants of dihedral angle potential energy terms describing rotation of the methyl groups, *k*_*dih*_, by Hoffmann et al.^29^ and starting points for the parameter search in this study

**Table 2:** Methyl-group content in the studied systems.^*a*^

| System | $C^\beta$ | $C^\gamma$ | $C^\delta$ | $C^\epsilon$ |
| --- | --- | --- | --- | --- |
| T4L | 15/16 | 24/40 (21/38) | 30/40 | 4/5 |
| CI2 | 3/3 | 28/29 (28/29) | 13/16 | 0/2 |
| Ubiquitin | 2/2 | 13/22 (10/15) | 16/25 | 1/1 |

<sup>a</sup> Numbers before and after the slash show measured/total number of methyl groups, respectively. Numbers in parentheses show $C^\gamma$ methyl groups excluding THR.

**Table 3:** Methyl-group-specific changes, Δ*k*_*dih*_(in kJ/mol), to the force constants of dihedral angle potential energy terms used for the parameter search in this study. *i* = 0 corresponds to the HMS force field by Hoffmann et al^29^

| $i$ | $C^\beta$ | $C^\gamma$ | $C^\delta$ | $C^\epsilon$ |
| --- | --- | --- | --- | --- |
| -2 | -0.07544 | -0.32738 | -0.17625 | -0.33995 |
| -1 | -0.07254 | -0.31479 | -0.16947 | -0.32687 |
| 0 | -0.06964 | -0.30220 | -0.16270 | -0.31380 |
| 1 | -0.06673 | -0.28960 | -0.15592 | -0.30072 |
| 2 | -0.06383 | -0.27701 | -0.14914 | -0.28765 |
| 3 | -0.06093 | -0.26442 | -0.14236 | -0.27457 |
| 4 | -0.05803 | -0.25183 | -0.13558 | -0.26150 |
| 5 | -0.05513 | -0.23924 | -0.12880 | -0.24842 |
| 6 | -0.05223 | -0.22665 | -0.12202 | -0.23535 |
| 7 | -0.04932 | -0.21405 | -0.11524 | -0.22227 |
| 8 | -0.04642 | -0.20146 | -0.10846 | -0.20920 |
| 9 | -0.04352 | -0.18887 | -0.10168 | -0.19612 |
| 10 | -0.04062 | -0.17628 | -0.09490 | -0.18305 |
| 11 | -0.03772 | -0.16369 | -0.08812 | -0.16997 |

**Table 4:** Overview of the simulations and force fields used in this study. *i* = 0 corresponds to the HMS force field by Hoffmann et al.^29^ a99SB*-ILDN combines several different force field modifications^38,39,44,60^

| Force field | Simulations |
| --- | --- |
| $i=-2$ | 5x1 μs T4L |
| $i=-1$ | 5x1 μs T4L |
| $i=0$ | 5x1 μs T4L, 5x1 μs CI2, 5x1 μs ubiquitin |
| $i=1$ | 5x1 μs T4L |
| $i=2$ | 5x1 μs T4L |
| $i=3$ | 5x1 μs T4L |
| $i=4$ | 5x1 μs T4L |
| $i=5$ | 5x1 μs T4L |
| $i=6$ | 5x1 μs T4L |
| $i=7$ | 5x1 μs T4L |
| $i=8$ | 5x1 μs T4L |
| $i=9$ | 5x1 μs T4L |
| $i=10$ | 5x1 μs T4L |
| $i=11$ | 5x1 μs T4L |
| $C_{\text{corr.}}^{\gamma}$ | 5x1 μs T4L, 5x1 μs CI2, 5x1 μs ubiquitin |
| a99SB*-ILDN | 5x1 μs T4L |

Changing *k*_*dih*_ results in different behaviours for the different types of methyl groups (Fig. 1 A-D): Increasing *k*_*dih*_ for C^*β*^ towards the original force field systematically worsens the agreement with the experimental data. C^*γ*^ shows the most distinct changes and the 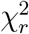 decreases with increasing *k*_*dih*_ until a minimum is reached around *i* = 9. At this point, 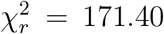 is reached which is almost half of the initial 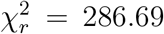 from the force field reparameterised by Hoffmann et al. In contrast, the 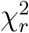 for C^*δ*^ methyl groups appear to remain mainly unchanged (note the outlier of one individual simulation at *i* = 4). Finally, the agreement between experimental and back-calculated relaxation rates of C^*ϵ*^ groups seems to get slightly better with increasing Δ*k*_*dih*_, however, the 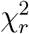’s are somewhat noisy which might be due to the small number of C^*ϵ*^ methyl groups in T4L (we analysed 4/5 C^*ϵ*^ groups) and thus there might be a risk of over-fitting.

**Figure 1:**
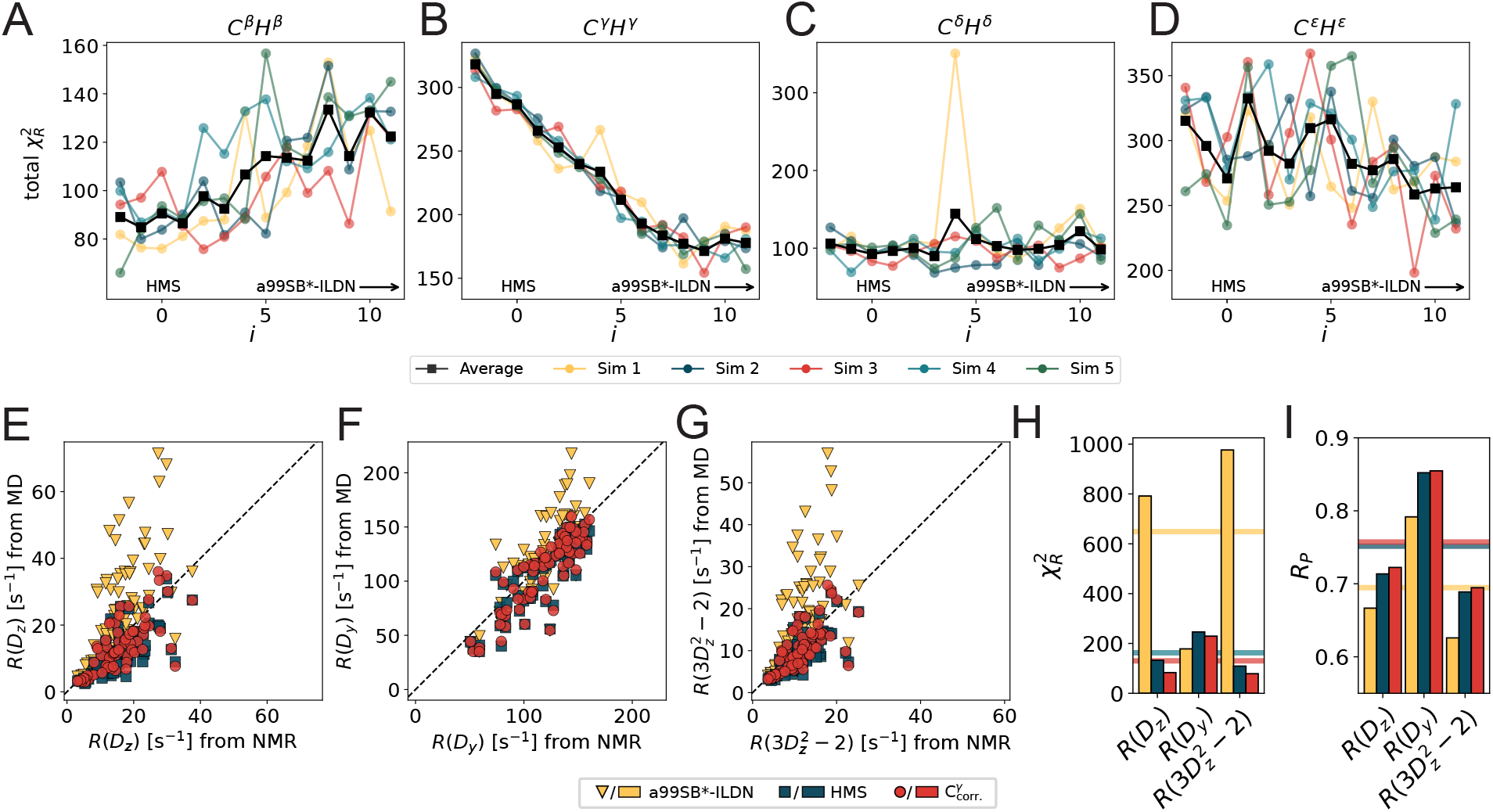
Parameterisation of potential energy barriers of methyl group rotation using T4L. (A-D) total 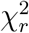 (average of 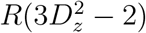 and *R*(3*D*^2^ *−* 2)) of different methyl groups as a function of Δ*k*. (E-G) *R*(*D*_*z*_), *R*(*D*_*y*_) and 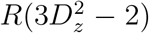, respectively, from the original force field^38,39,44,60^ (yellow triangles), the HMS^29^ (blue squares) and the 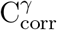 (redcircles) optimised force fields compared to experimental data. (H-I) 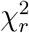 and Pearson correlation coefficient of the data shown in E-G. Horizontal lines show corresponding averages over the three relaxation rates.

Based on this data, we decided to modify only the methyl rotation barriers of C^*γ*^*H*^*γ*^-bearing residues for two reasons: First, our data suggest that *k*_*dih*_ is already near an optimum for C^*β*^, C^*δ*^ and C^*ϵ*^ while we see stark improvements for C^*γ*^; Second, our previous work^34^ also showed that mainly VAL and ILE are problematic which are both C^*γ*^*H*^*γ*^-bearing amino acids. For these reasons, we decided to use *i* = 9, corresponding to a Δ*k*_*dih*_ of *−*0.188 87 kJ*/*mol. We tested if reparameterising one type of methyl group influences the dynamics of the other methyl groups, and we see no substantial difference on those residues upon modifying the C^*γ*^ rotation barriers (Fig. S1-3). From this, we conclude that the methyl-spinning of the individual methyl groups is largely uncoupled from each other and thus *k*_*dih*_ can be modified individually.

### 2.2 Testing the modified force-field

As expected from the results described above, simulations of T4L performed with our forcefield modification applied to C^*γ*^*H*^*γ*^ agree better with experiments than both the original and the force-field by Hoffmann et al (Fig. 1 E-I). The biggest relative improvements are observed for *R*(*D*_*z*_) and 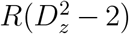 in both reduction of the 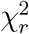 (Fig. 1 H) and increase of the Pearson correlation coefficients (Fig. 1 I); *R*(*D*_*y*_) improved less, presumably due to its dependence on *J*(0). These improvements, however, are expected because the parameter scan and the resulting selection of *k*_*dih*_ was based on simulations of T4L. To test the new force-constant, we used two independent protein systems, namely CI2 and Ubiquitin. The improvement of the force-field becomes specifically apparent in a test system like CI2 containing where 29 out of 50 methyl groups are of the C^*γ*^ type (Table. 2). For CI2, we calculated the deuterium relaxation rates *R*(*D*_*z*_), *R*(*D*_*y*_), 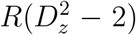, and, in addition, 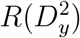 (Eq. 5-8) from five 1 µs-long simulations using the force-field by Hoffmann et al., and our optimised version and compared them to previously measured NMR relaxation data (Fig. 2).^11^ The Pearson correlation coefficient improves substantially in the case of all four relaxation rates with respect to the previously optimised force field by Hoffmann et al (Δ*R*_*P*_ = 0.16, Δ*R*_*P*_ = 0.06, Δ*R*_*P*_ = 0.11, Δ*R*_*P*_ = 0.18 for *R*(*D*_*z*_), *R*(*D*_*y*_), 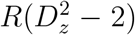 and 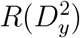, respectively; Fig. 2 A-D). This improvement is driven by the increased agreement between experimental and simulated C^*γ*^*H*^*γ*^ relaxation rates, and is also reflected in the halving of the 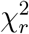. We note a very small increase in the 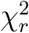 for the remaining methyl groups (Fig. 2 E), but also that this change is much smaller relative to the improvement of the C^*γ*^*H*^*γ*^ relaxation rates. This is also apparent from Fig. S2 which shows that the Pearson correlation coefficient of the calculated relaxation rates from unmodified methyl groups between the two force field is nearly 1.

**Figure 2:**
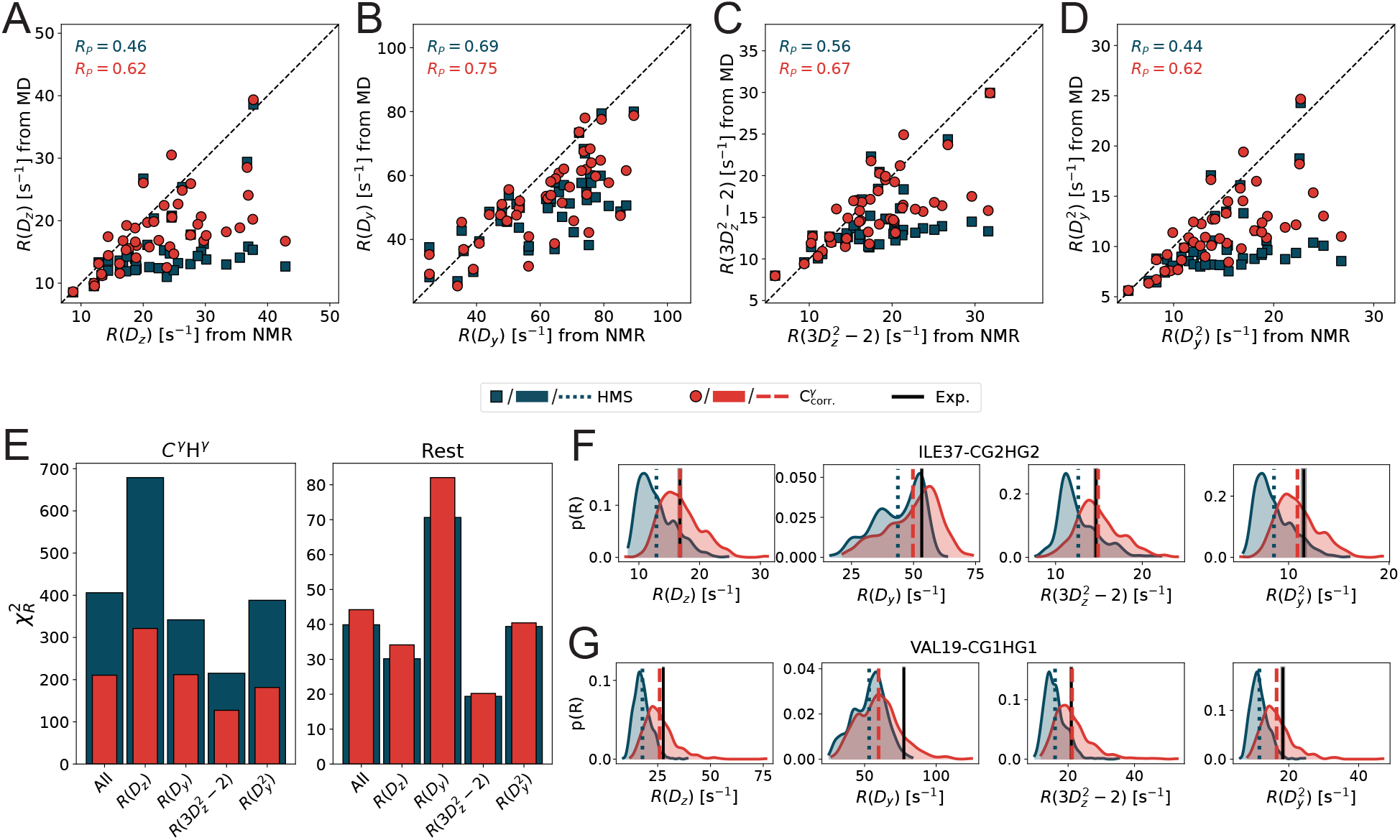
Testing of the optimised 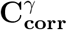 force field on CI2 relaxation data. (A-D) Comparison of the HMS (blue squares) and the 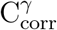 (red circles) force fields to experimental NMR relaxation data. (E) Agreement as shown by 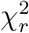 between experimental relaxation rates and the HMS and 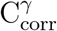(blue and red bars, respectively) force fields for C^*γ*^H^*γ*^ alone (*left panel*) and the remaining methyl groups (*right panel*). (F-G) Representative distributions of relaxation rates from the HMS (blue densities) and 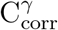 (red densities) force-fields of ILE37-C^*γ*2^H^*γ*2^ (F) and VAL19-C^*γ*1^H^*γ*1^ (G) and their respective averages (dotted and dashed vertical lines). Experimental values and uncertainties are shown as black lines and shading, respectively.

The modification of the force constant affecting methyl spinning also affects the distributions of relaxation rates from the individual trajectory blocks which are used to calculate average relaxation rates from the simulations (see methods section). As an example, we show the distributions of each relation rate for the C^*γ*^ methyls of ILE37 (Fig. 2F) and VAL19 (Fig. 2G) and their corresponding averages which are compared to experimental data. In all cases, there is improved agreement between the calculated and measured relaxation rates. The accuracy is strongly improved considering that in many cases there was almost no overlap between the distribution of relaxation rates of the individual simulation blocks and the experimentally-measured average rate. We also note that the distributions become slightly broader (Fig. S4B).

Next, we tested our force field modification on ubiquitin, a protein that is commonly used for both NMR relaxation studies and benchmarking force fields. ^22,33,38,39,61,62^ We calculated a set of five relaxation rates including *R*(*D*_*z*_), *R*(*D*_*y*_), 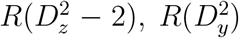 and *R*(*D*_*y*_*D*_*z*_ + *D*_*z*_*D*_*y*_) (Eq. 5-9) from five 1 µs-long simulations and compared them to NMR relaxation data by Liao et al.^61^ (Fig. 3). Although ubiquitin has a lower C^*γ*^ content than CI2, we observe overall improvements of the agreement between experimental and calculated relaxation rates (Table 2). Fig. 3 A-E shows that the Pearson correlation coefficient increased most for *R*(*D*_*z*_), 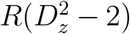 and 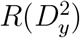 (by 0.21, 0.21 and 0.26, respectively) whereas it remained almost the same for *R*(*D*_*y*_) (Δ*R*_*P*_ = 0.01)and *R*(*D*_*y*_*D*_*z*_ + *D*_*z*_*D*_*y*_) (Δ*R*_*P*_ = − 0.01). Interestingly, the initial agreement is already quite good for the latter two relaxation rates with *R*_*P*_ = 0.77 and *R*_*P*_ = 0.79. 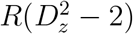 and *R*(*D*_*y*_*D*_*z*_ + *D*_*z*_*D*_*y*_) are the only two relaxation rates which also depend on spectral densities at the zero frequency and thus are also more influenced by the tumbling rate *τ*_*R*_. To make it easier to compare the different force fields and simulations, we used the same experimentally derived *τ*_*R*_ in the calculations of relaxation rates from the different force fields which partly explains why those two relaxation rates are less influenced by the force field modification. Similar to CI2, we also see an improvement of the agreement between experimental and calculated C^*γ*^*H*^*γ*^ relaxation rates when using our modified force field (Fig. 3F). At the same the 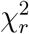 stays the same or slightly increases for the remaining methyls. Looking at the distributions of the relaxation rates from the MD simulations we also see a slight broadening effect which also results in a better overlap with the experimental measurements (Fig. 3G-H and Fig. S4C).

**Figure 3:**
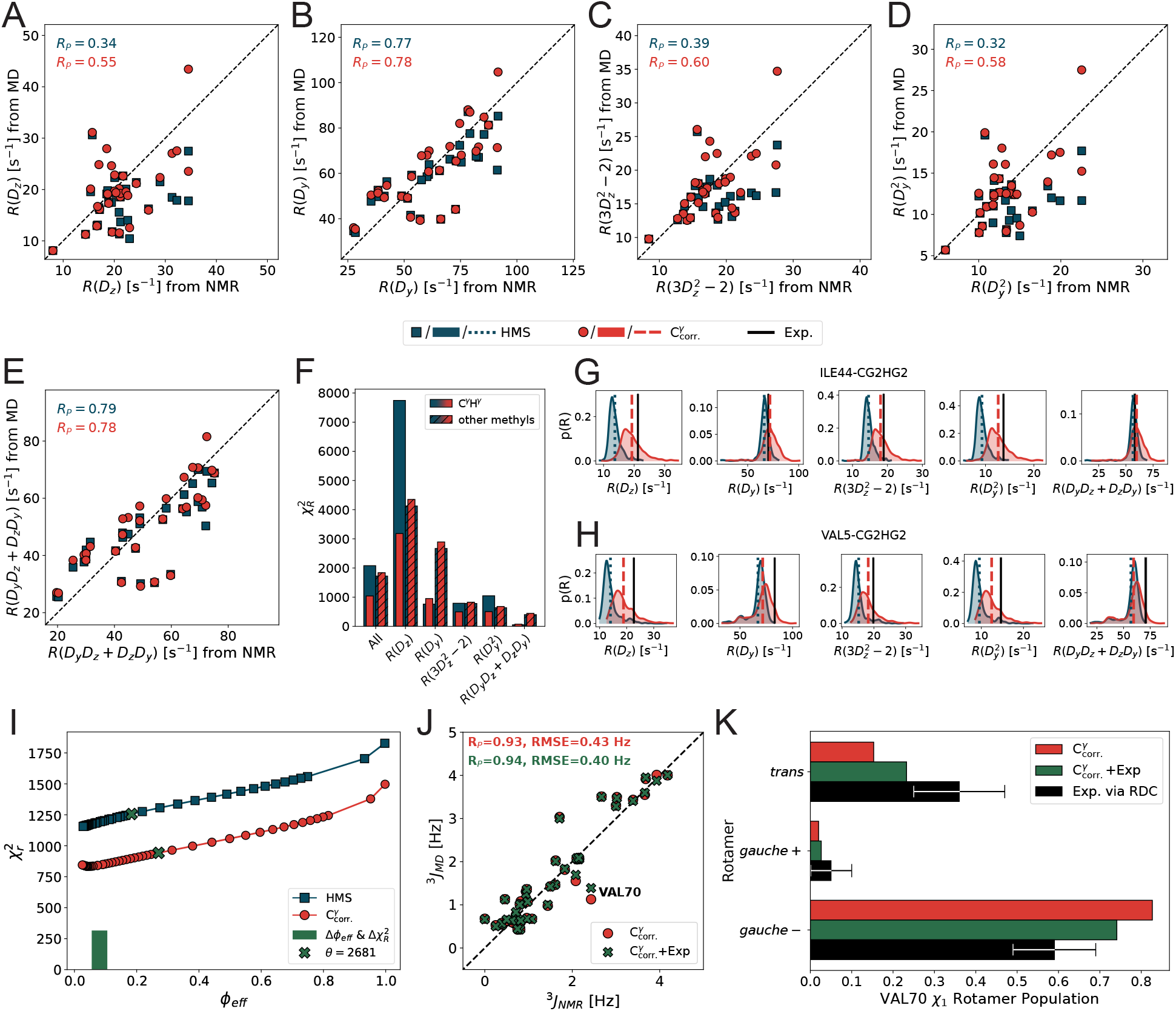
Testing of the optimised 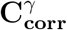 force field on ubiquitin relaxation data. (A-E) Comparison of the HMS (blue squares) and the 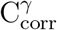 red circles) force fields to experimental NMR relaxation data by Liao et al.^61^ (F) 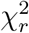 reporting on the agreement between experimental relaxation rates and rates from simulations with the HMS and 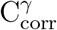 (blue and red bars, respectively) force fields for C^*γ*^H^*γ*^ alone (open bars) and the remaining methyl groups (hatched bars). (G-H) Representative distributions of C^*γ*2^H^*γ*2^ relaxation rates from the HMS (blue densities) and 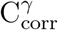 (red densities) force-fields of ILE44 (G) and VAL5 (H) and their respective averages (dotted and dashed vertical lines). Experimental values and uncertainties are shown as black lines, respectively. (I) Behaviour of the reduced *χ*^2^ vs *ϕ*_eff_ during the reweighting with ABSURDer for the HMS (blue squares) and the 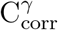 (red circles) force fields using different values of *θ*. The green cross shows the respective points for *θ* = 2681 determined using *k*-fold cross validation and the bar of the same colour represents the difference between the two force fields. (J) Cross-validation with C’/N-C^*γ* 3^J scalar couplings^62^ before (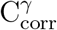, red circles) and after (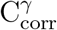 +Exp., green crosses) reweighting the 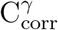 simulations against relaxation rates. (K) *χ*_1_ rotamer populations of VAL70 before (red bars) and after (green bars) reweighting. Experimental values including standard deviation (black bars) are derived from side-chain residual dipolar couplings. ^62^

### 2.3 Reweighting MD trajectories

We were interested to examine if we could further improve the agreement between the experimental and calculated relaxation rates and thus applied our previously described reweighting method, ABSURDer (average block selection using relaxation data with entropy restraints).^34^ In ABSURDer, the simulations are fitted to experimental data by minimising the functional *𝒯* = *χ*^2^(**w**) *− θS*(**w**), where *χ*^2^ quantifies the agreement between experiments and simulations, *S* is a measure of the extent of reweighting and **w** are the fitting parameters (weights of each simulation block). This is achieved by fitting **w** to minimize *𝒯* (see Eq. 11 and methods for additional details). In order to balance how tight the experimental data are fitted (through minimisation of *χ*^2^(**w**)) and how much of the original simulation is retained (regularised by the *S*(**w**) entropy term), one has to select an adequate value for *θ*. The extent of reweighting can be judged by the parameter *ϕ*_eff_ = exp(*S*), which provides a measure of the effective fraction of blocks retained for a given set of weights and thus gives information on the amount of simulation data that needs to be upweighted to match a given set of experimental restraints. We compared the behaviour of 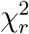 and *ϕ*_eff_ using different values of θ for when reweighting simulations generated using either the HMS of 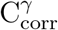 force fields (Fig. 3I). We highlight three observations from Fig. 3I: First, before reweighting the 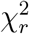 (*prior*) for the 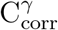 force field is lower than that for the HMS force field, in line with the improved agreement shown in Figure 3F. Second, the relative improvement of the new force field after reweighting with a selected *θ* is better than the *posterior* of the force field by Hoffmann et al (rel. 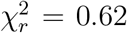 and rel. 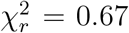, respectively, where rel. 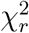 refer to the relative decrease of 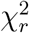 from reweighting) indicating that, here, reweighting is more effective. Third, more simulation data is retained after reweighting with the given *θ* = 2681 using 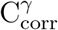 (*ϕ*_eff_ = 0.27) compared to the HMS force field (*ϕ*_eff_ = 0.18).

One feature of reweighting is that the optimised weights can be used to calculate other observables which might in favourable cases lead to better agreement with other experimental data compared to using the unrefined simulations. To test this, we calculated C’/N-C^*γ* 3^*J*-couplings using both the initial, uniform weights and the optimised weights and compared them to experimental values reported by Chou et al.^62^ (Fig. 3J). We note that the agreement of the ^3^*J*-couplings from the *prior* is already high (Pearson correlation of 0.93 and RMSE=0.43 Hz) which also relates to the fact that the original force field was initially validated against these and other scalar couplings.^39^ Nonetheless, we see further, although small, improvements upon reweighting the simulations against relaxation data. Intriguingly, we find the biggest improvement for VAL70. It was previously shown, that VAL70 undergoes slow motions that are linked to rotameric jumps,^62,63^ and previous work showed imperfect agreement between simulations and experiments for this residue. ^29^ We find that the *χ*_1_ rotamer distribution of VAL70 changes upon reweighting and the increase of the *trans* population with the coupled decrease of the *gauche-* rotamer population is also in better agreement with experimentally obtained rotamer populations by Chou et al ^62^ (Fig. 3K). Altogether, this demonstrates that reweighting can be used to further refine the simulations and that fitting against relaxation data also improves the agreement of equilibrium observables such as dihedral angle distributions. We also note that since ABSURDer takes into account both the amplitudes and timescales of motion, the reweighting can appropriately deal with the fact that relaxation data and scalar couplings are affected by the motions in different ways.^62,63^

## 3 Conclusions

Here, we have shown that we can further improve the ability of MD simulations to capture side chain dynamics as probed by NMR relaxation. In particular, we have presented an approach to fit torsional potential energy terms to NMR relaxation data by directly calculating the relaxation parameters from MD simulations. We performed this optimisation targeting data in T4L and demonstrate the improvement using CI2 and ubiquitin. Furthermore, we present an example of how the optimised *prior* also enables a more efficient application of *a posteriori* reweighting of the MD trajectory using ABSURDer.

However, our data also highlights that there is still some room for improvements in the prediction of NMR relaxation parameters which could have various reasons. Although the mechanisms of spin relaxation are well understood and the use of a spectral-density-based approach to calculate NMR relaxation parameters is less dependent on assumptions compared to model-free approaches,^64–67^ there might still be some effects that are not fully captured by the applied model to calculate relaxation rates^20^ such as the treatment of overall tumbling or the limited number of exponentials in the fitting of the time correlation functions. For further discussion in this regard we point to our previous work on this subject.^34^ We note also that there might be small effects resulting from deuteration that are not captured by the simulations.^68–71^ Finally, there could be remaining force field related issues. This could, for instance, relate to insufficient training data and having access to more protein systems with available deuterium spin relaxation data could be useful. Other types of force field parameters can potentially influence the methyl group dynamics such as non-bondend terms describing the interactions of the probed side chains with their chemical environment. Exploring this parameter space, however, would require more extensive approaches for a wider scan of parameters and thus also more simulations. One strategy could be the use of reweighting techniques in the force field fitting procedure that would then lower the required amount of simulations,^36,37,45,72–74^ though these would in need to be adapted to time-dependent experimental data.^75^ Other possibilities include combining additional QM calculations with optimisation against the experimental data.

In summary, we present a modified force field that gives rise to improved agreement between simulations and side chain NMR relaxation experiments. We envisage that the force field will enable more detailed assessments of the motions probed by NMR relaxation experiments, and focus future force field developments on other aspects of the force field that might require improvements.

## 4 Methods

### 4.1 Parameter scanning

To find an improved force constant for methyl spinning we set up a parameter search using the force field modification by Hoffmann et. al as a starting point.^29^ Next, we perform a stepwise increase of the force constants of the dihedral angle potential energy terms underlying the rotation of the methyl (*k*_*dih*_). The modifications (Δ*k*_*dih*_) by Hoffmann et al. are reported in Table 1. We apply the same relative increase to each methyl group (except for the C^*γ*^ of THR) and set 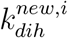 in the following way:

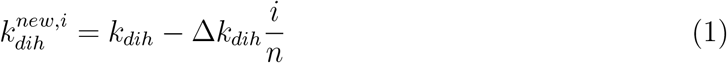

where *n* = 24 and *i* ranges from -2 to 11 in steps of 1 (Table 3). For each step (i.e parameter set), we run five 1 µs-long simulations of T4L (Table 4), calculate the deuterium relaxation rates *R*(*D*_*z*_), *R*(*D*_*y*_)and 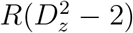 and compare them to experimental measurements obtained at magnetic field strength corresponding to 950 Hz. ^30^

### 4.2 MD simulations

We used the X-ray structure of the cysteine-free T4L Ser44Gly mutant (PDB 107L) ^76^ where we mutated GLY44 back to a serine as the starting configuration for T4L. We also used an X-ray structure for CI2 (PDB 7A1H) ^77^ and a randomly selected model (no. 60) from an NMR structure (PDB ID 1XQQ) of ubiquitin.^78^ We parameterised the systems using the AMBER ff99SB*-ILDN/TIP4P-2005 protein force field^38,39,44,60^ in combination with the modifications to the methyl rotation barriers by Hoffmann et al.^29^ and herein (see below for details).

All MD simulations were carried out with GROMACS v2020. ^79^ Simulations were set up with a 1.2 nm distance between the proteins and the cell borders and used a periodic trun-cated dodecahedron of 400, 200 and 176 nm^3^ volume as simulation boxes for T4L, ubiquitin and CI2, respectively. We solvated the system with 12303 (T4L), 6107 (ubiquitin) and 5418 (CI2) TIP4P-2005 water molecules,^60^ retaining crystal waters if present. We neutralised the systems with Na^+^ and Cl^*−*^ ions, further matching experimental ion concentrations if necessary. The systems were minimized with 50000 steps of the steepest descent and equilibrated for 200 ps in the NPT ensemble, using harmonic position restraints (force constants of 1000 kJmol^*−*1^nm^*−*2^) on the heavy atoms of the protein. We used the leap-frog algorithm to integrate the equations of motion with a time-step of 2 fs. We applied a cutoff of 1 nm for the non-bonded interactions and employed Particle-mesh Ewald summation for long-range electrostatics^80^ with the 4th-order cubic interpolation and 0.16 nm Fourier grid spacing. We ran the production simulations in the NPT ensemble, using the velocity rescaling thermostat^81^ with a reference temperature of 298 K and thermostat timescale of *τ* = 1 ps and the Parrinello–Rahman barostat^82^ with a reference pressure of 1 bar, a barostat timescale *τ* = 2 ps, and an isothermal compressibility of 4.5 *×* 10^*−*5^ bar^*−*1^. For each simulation system, we performed a set of five 1 µs production runs (Table 4). Finally, we saved the protein coordi-nates every 1 ps and, after running the simulations, we removed the overall tumbling of the protein by fitting to a reference structure.

### 4.3 Calculation of NMR parameters from MD simulations

We used spectral density mapping to calculate NMR relaxation rates as previously described.^30,34^ We started by splitting each simulation into non-overlapping, 10 ns long blocks resulting in a total of 500 blocks per system and parameter set. Next, we calculated the internal time correlation function (TCF), *i*.*e*. after removing global tumbling, of the three 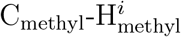 (*i* = 1, 2, 3) bond vectors up to a maximum lag-time of 5 ns for all side-chain methyl groups. The internal TCFs were then fitted with six exponential functions and an offset,

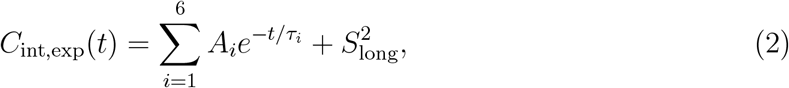

where 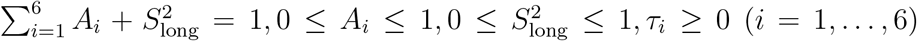 and 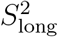 is the long-time limit order parameter; however, no physical meaning should be attributed to these amplitudes and timescales.^16^ Previous work showed that an axially symmetric tumbling model describes T4L, which is why we introduced methyl-specific global tumbling times *τ*_R,*i*_ by multiplying the internal TCF with a single-exponential, thus yielding the total TCF:^30^

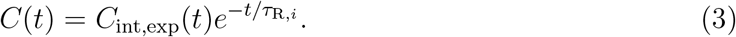

We applied an isotropic tumbling model to the other two systems using experimental values of *τ*_R,*iso*_ = 5.00 ns and *τ*_R,*iso*_ = 3.99 ns for ubiquitin^61^ and CI2,^11^ respectively.

Afterwards, we performed a Fourier transformation of the total TCF to obtain a spectral density function:

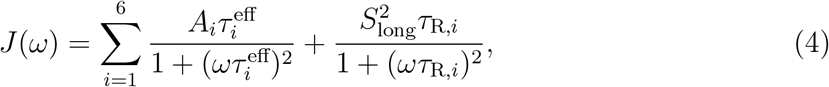

where 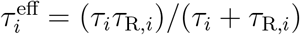. Finally, we calculated up to five relaxation rates from *J*(0), *J*(*ω*_D_), and *J*(2*ω*_D_) via

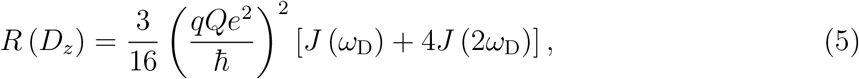

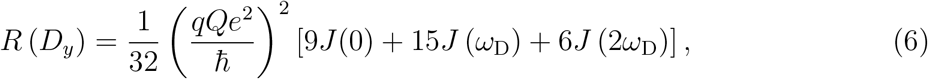

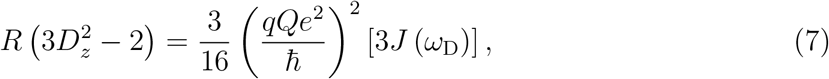

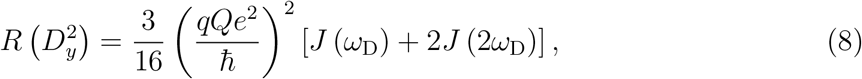

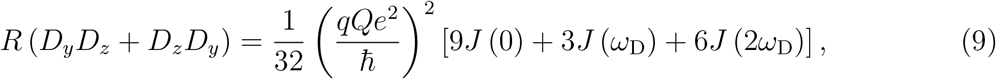

where (*qQe*^2^*/*ħ)^2^ is the quadrupolar coupling constant of deuterium. We used 145.851 MHz, 92.124 MHz and 115.150 MHz as the Larmor frequency *ω*_D_ of deuterium for T4L, ubiquitin and CI2, respectively, matching Bruker magnetic field strengths of 22.3160 T, 14.0954 T and 17.6185 T at which the experimental NMR relaxation rates were measured. In this way, we calculated up to five relaxation rates for each methyl group and each of the 500 blocks per simulation system, which were then averaged over the blocks and directly compared to experimental data. For reweighting, we used the blocks prior to averaging (see below).

To calculate C’-C^*γ*^ and N-C^*γ* 3^*J*-couplings of VAL, ILE and THR residues, we used the generalised Karplus expression:^83^

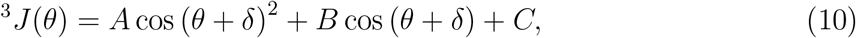

where *θ* is the relevant torsion angle, and *δ* is a small phase shift. We used the corresponding coefficients *A, B, C* and *δ* as parameterised by Chou and co-workers.^62^

### 4.4 Reweighting with ABSURDer

We use our previously described reweighting method ABSURDer^34^ to further optimise the agreement of our ubiquitin simulations with experimental NMR measurements. To do so, we find an optimised set of weights, **w**, associated with the blocks from the simulations by minimising the functional:

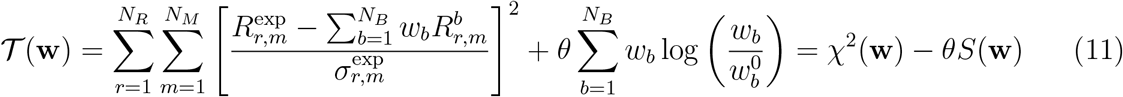

where *r* runs over the *N*_*R*_ types of experimental NMR relaxation rates, *N*_*M*_ is the overall number of measured rates of type *r, N*_*B*_ represents the number of blocks which the trajectory has been cut into, 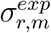 is the experimental error on the *m*-th rate of the *r* -th type and constraining Σ*w*_*b*_ = 1. The second term in the functional represents a regularisation in the form of a Shannon relative-entropy,^73,84,85^ *S*(**w**), preventing over-fitting when the new weights (*w*_*b*_) deviate too much from the initial set of weights 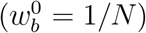. The parameter *θ* is used to balance the contribution from the *prior* (through the *S*(**w**)) and the experiments (given by the *χ*^2^). We used *k*-fold cross-validation to choose *θ* from a range of values as previously described.^11^ Briefly, we split the relaxation data into training (80%) and test (20%) sets, and reweighted the training set against experimental data using a range of different *θ* values. The obtained weights were then applied to the test set and the agreement of both was measured as a relative 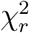. We repeated this procedure *k* = 10 times and selected *θ* according to the minimum of the relative 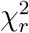 in the test set (Fig. S5). The parameter *ϕ*_eff_ = exp *S*(**w**), can be used as a measure of the effective fraction of blocks retained for a given set of weights.

### 4.5 Data and code availability

Data and code generated for this work is available via https://github.com/KULL-Centre/_2023_Kummerer_nmrFF/. ABSURDer is available at https://github.com/KULL-Centre/ABSURDer

## 5 Acknowledgments

We are grateful to Profs. Lars V. Schäfer and Kaare Teilum for discussions about our work. We acknowledge support by a grant from the Lundbeck Foundation to the BRAINSTRUC structural biology initiative (155-2015-2666, to K.L.-L.). We acknowledge access to computational resources from the ROBUST Resource for Biomolecular Simulations (supported by the Novo Nordisk Foundation; NNF18OC0032608), the Danish National Supercomputer for Life Sciences (Computerome), and the Biocomputing Core Facility at the Department of Biology, University of Copenhagen.

## Notes

https://github.com/KULL-Centre/_2023_Kummerer_nmrFF/

### Competing Interest Statement

The authors have declared no competing interest.

