## Supporting Figures for "Fitting Force Field parameters to NMR Relaxation Data"

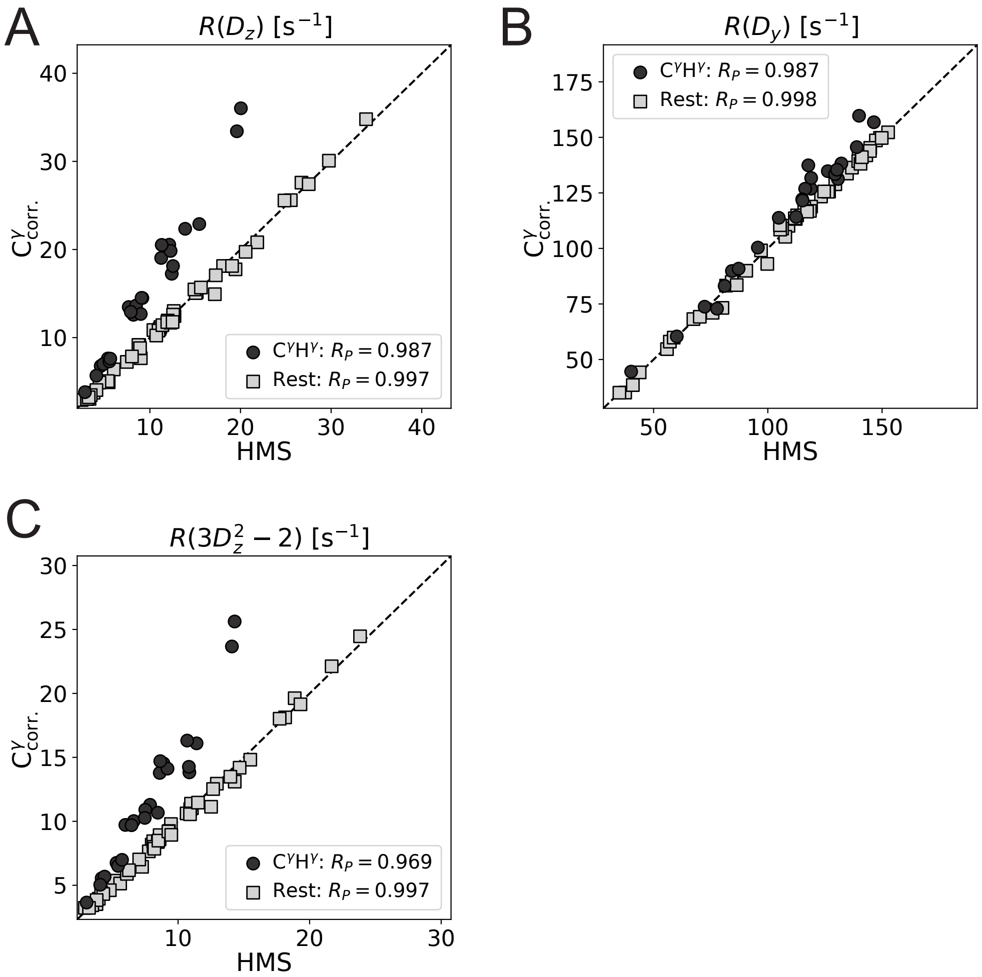


Figure S1: **Independence of methyl group dynamics in T4l.** Scatter plots comparing (A) $R(D_z)$ , (B) $R(D_y)$ , and (C) $R(D_z^2 - 2)$ of T4L from the C$^{\gamma}_{\text{corr.}}$ force field and the HMS force field. Relaxation rates from $C^{\gamma}H^{\gamma}$ methyl groups are coloured in anthrazite, others are coloured in light grey.

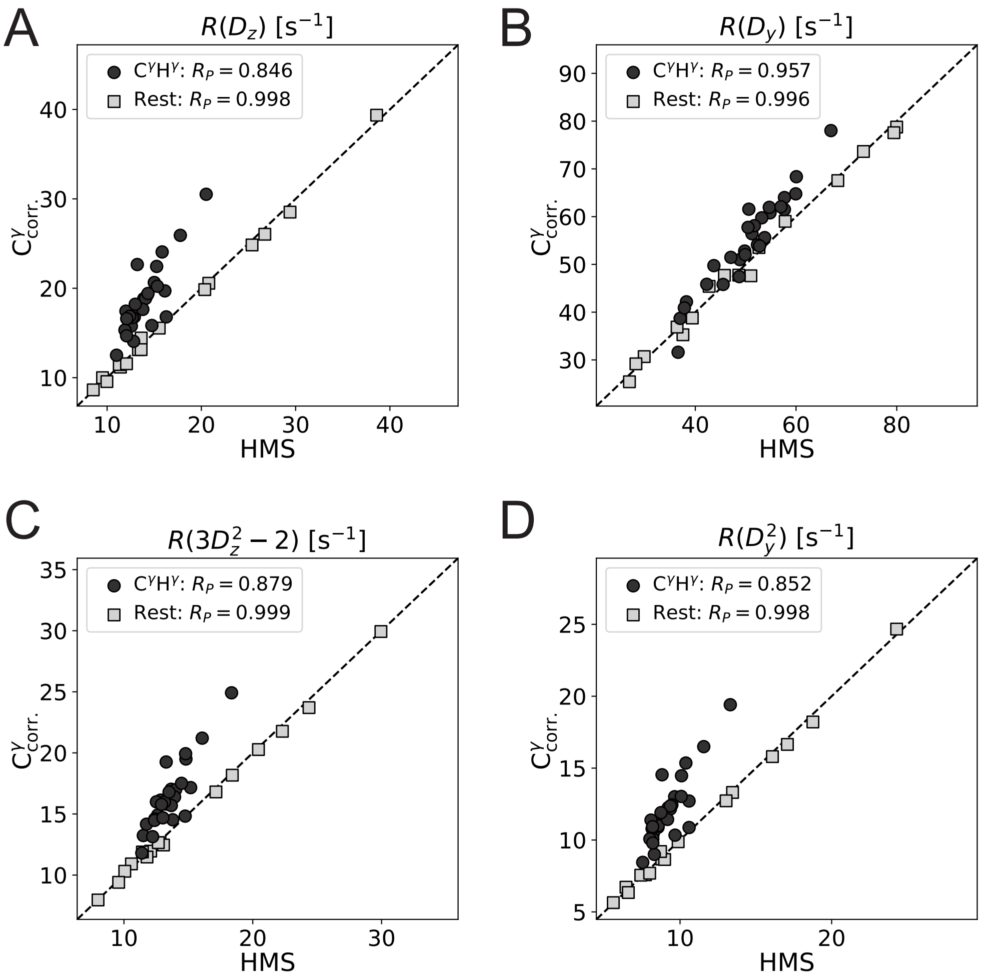


Figure S2: **Independence of methyl group dynamics in CI2.** Scatter plots comparing (A) $R(D_z)$ , (B) $R(D_y)$ ,(C) $R(D_z^2-2)$ , and (D) $R(D_y^2)$ of CI2 from the $C^{\gamma}_{corr.}$ force field and the HMS force field. Relaxation rates from $C^{\gamma}H^{\gamma}$ methyl groups are coloured in anthrazite, others are coloured in light grey.

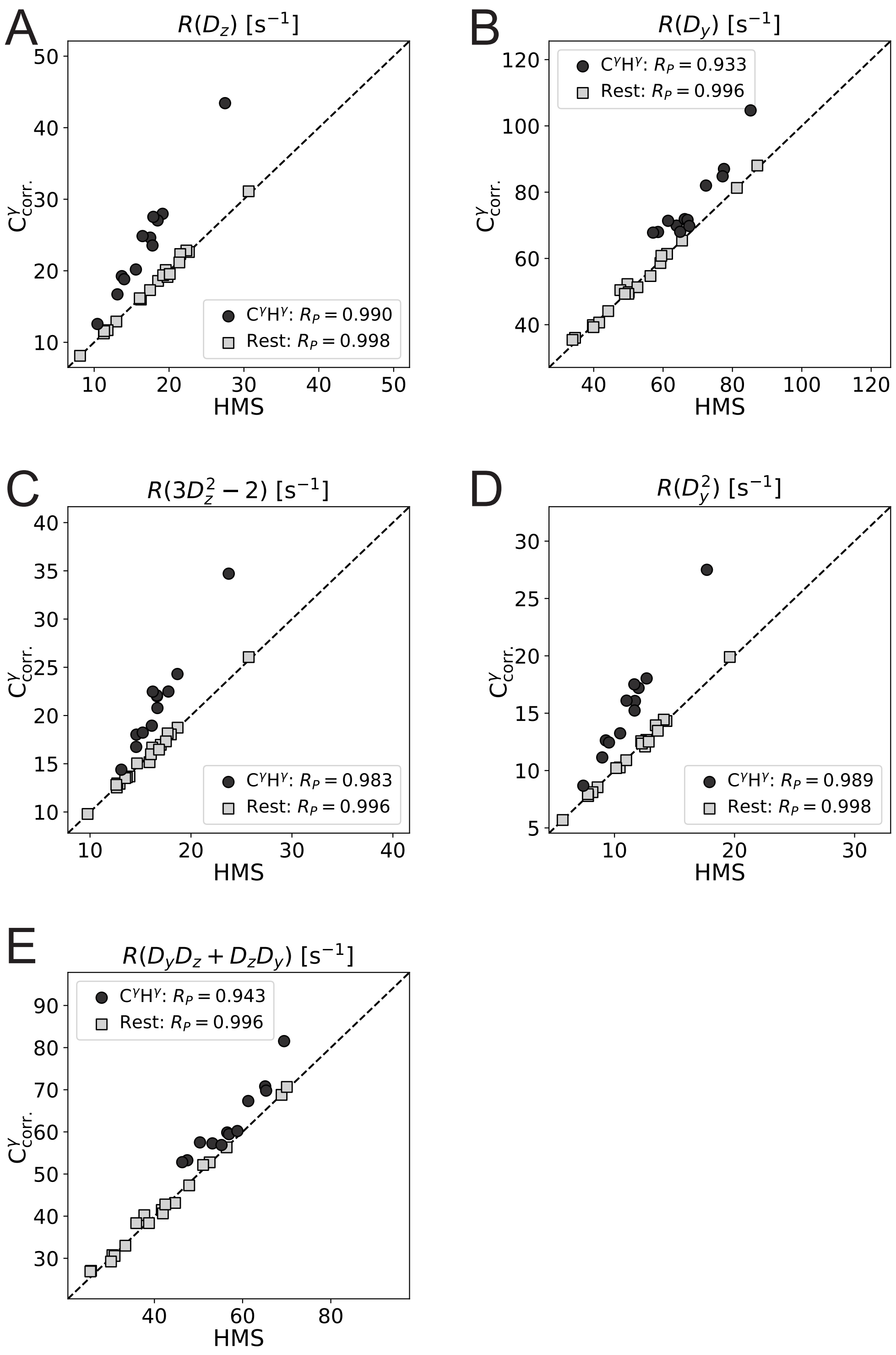


Figure S3: **Independence of methyl group dynamics in ubiquitin.** Scatter plots comparing (A) $R(D_z)$ , (B) $R(D_y)$ , (C) $R(D_z^2 - 2)$ (D) $R(D_y^2)$ , and (E) $R(D_yD_z + D_zD_y)$ of Ubiquitin from the $C^{\gamma}_{\text{corr.}}$ force field and the HMS force field. Relaxation rates from $C^{\gamma}H^{\gamma}$ methyl groups are coloured in anthrazite, others are coloured in light grey.

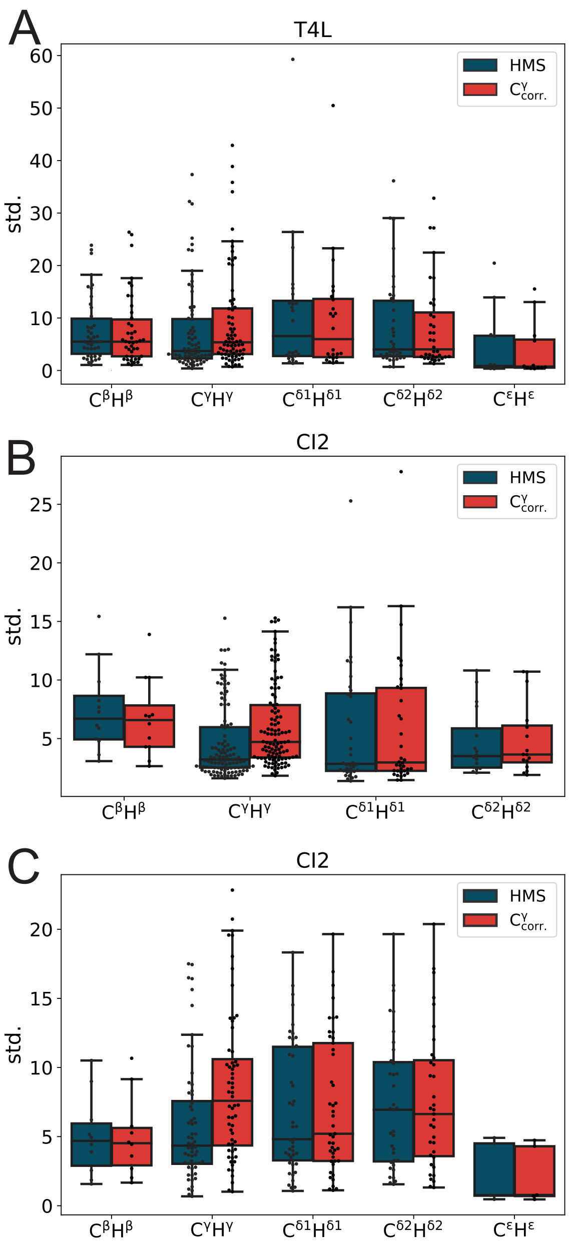


Figure S4: **Standard deviation of relaxation rate distributions.** (A-C) Comparison of the methyl-group-specific standard deviation of all relaxation rates over the trajectory blocks from simulations using the HMS (blue) and the $C^{\gamma}_{corr.}$(red) force fields.

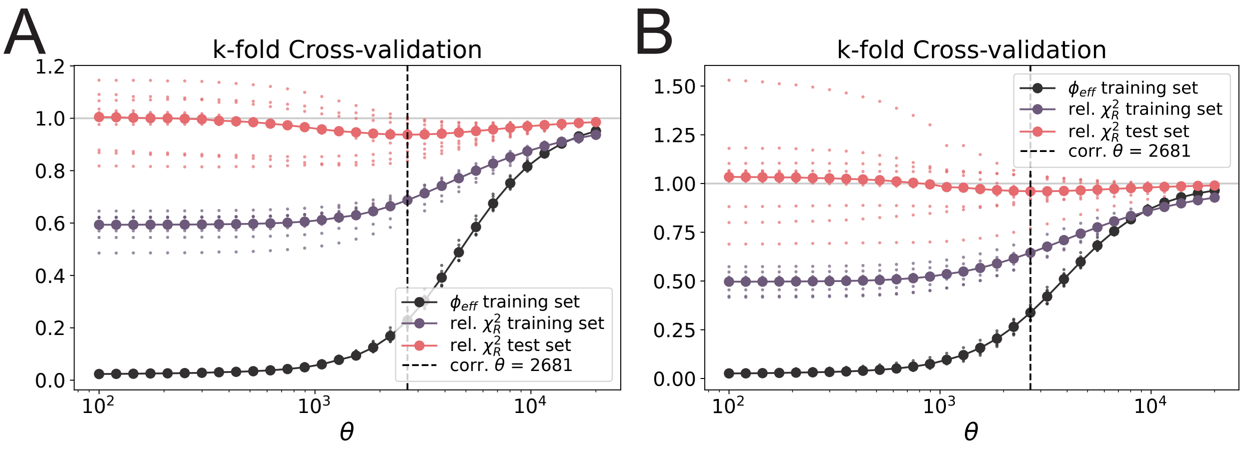


Figure S5: **Cross-validation for the selection of $\theta$.** Relative $\chi^2$ of simulated relaxation rates from (A) the HMS and (B) the $C^{\gamma}_{corr.}$ force field to experimental NMR data of the training set (80% of the relaxation rates, black points), the test set (20% of the relaxation rates red points) and $\phi_{eff}$ as a function of $\theta$. The large points show the averages of the $k = 10$ individual cross-validations. The vertical lines shows the selected $\theta$.
